# Mapping the shared and unique structural asymmetry abnormalities of young children with autism and developmental delay/intellectual disability with normative models and their multimodal cascade

**DOI:** 10.1101/2023.12.10.571041

**Authors:** Shujie Geng, Yuan Dai, Edmund T. Rolls, Yuqi Liu, Yue Zhang, Lin Deng, Zilin Chen, Jianfeng Feng, Fei Li, Miao Cao

## Abstract

To understand the neural mechanism of autism spectrum disorder (ASD) concurrent with developmental delay/intellectual disability (DD/ID), it is essential to comprehensively take genetic, brain, and behavioural measurements as a whole and focus on subjects at early age. However, such research is still lacking.

Here, using the sMRI data of 1030 children under 8 years old, we employed developmental normative models to explore the atypical development of gray matter volume (GMV) asymmetry in individuals with ASD without DD/ID, ASD with DD/ID and DD/ID, and their associations with neurophysiological measures and transcription profiles.

By computing the individual deviations from typical controls, we observed an ASD-specific abnormal GMV laterality pattern that was more rightwards in the inferior parietal cortex and precentral cortex and noted abnormal within-group heterogeneity in the temporal pole. Specifically, ASD with DD/ID children exhibited more regional abnormalities; ASD without DD/ID children showed higher within-group variability; while children with DD/ID showed no significant abnormalities. However, there were no significant differences among the three groups. The GMV laterality of ASD without DD/ID children was associated with ASD symptoms, whereas that of ASD with DD/ID children was associated with both ASD symptoms and verbal IQ. Last, the GMV laterality of the ASD with DD/ID, ASD without DD/ID, and DD/ID groups was associated with shared and unique gene expression profiles, but the associations of the latter two groups with intellectual genes showed opposite effects.

Our findings illustrated the atypical development of regional structural lateralization in autistic children, which is associated with upstream genes and downstream behavioural performance. The differences and similarity between ASD and DD/ID additionally improve our standing to the neural mechanism of neurodevelopmental disorders comorbidity.

## Introduction

Autism spectrum disorder (ASD) is a typical neurodevelopmental disorder, that is difficult to effectively diagnose and treat due to its comorbidity with other neurodevelopmental disorders. It has now been recognized that atypical neural developments in the ASD population take place during the prenatal [1] to childhood stages [2–4], and symptoms appear before age 3 [5]. A substantial number of recent studies have investigated ASD-specific and shared pathological mechanisms with other neural developmental disorders in different modalities, including the genetics, transcriptomes, neural imaging and behavioural performance in adults. However, studies focused on young children and comprehensively exploring multimodal abnormalities are still lacking. Additionally, ASD and developmental delay/intellectual disability (DD/ID) co-occur with a high prevalence of 68% [6–8], and both are highly heritable, with genetic causes contributing to 30%-70% [9] and 25–50% of cases [10], respectively. The prevalence of DD/ID is estimated to be 1–3% in children under 5 years old [11]. Focusing on the multimodal cascade of brain measurements during the early-stage of life [12–16], referring to how upstream gene expression affects hierarchical brain structures and functions, ultimately leading to clinical manifestations, will be crucial for revealing the comorbidities and differences between ASD and DD/ID for understanding the neuropathological mechanisms of both disorders.

Notably, left-right asymmetry is a fundamental organizing feature of brain structure and function and becomes specialized with development [17]. Structural and functional brain asymmetry have been reported to be heritable [18, 19]; to be genetically modulated [20–22]; and are established early in neonates [23], preterm infants [24] and even fetuses [25, 26]. In typically developing individuals, asymmetry measurements of brain asymmetry, including morphological indexes [27–29], functional and structural connectivity [30, 31], and topological properties of functional and structural brain networks [32, 33], were linked with functions of language, perception and action, emotion and decision making [34]. Notably, those functions were commonly impaired in ASD and DD/ID patients, suggesting that brain asymmetry has great potential as an intermediate neural phenotype linking genetic factors and clinical manifestations for those atypical populations.

Previous neuroimaging research revealed atypical brain asymmetry in populations with ASD. Reduced leftwards structural and functional asymmetry in the frontal gyrus, superior temporal gyrus and other language-related regions were linked to language impairments [35–38]. Specifically, Postema et al. investigated structural brain asymmetry in ASD patients based on a large sample recruited from the ENIGMA consortium, with 1774 ASD patients and 1809 controls, and found mild alterations in cortical thickness asymmetry in the medial frontal, orbitofrontal, cingulate and inferior temporal areas; surface area asymmetry in the orbitofrontal cortex, and volume asymmetry in the putamen [39]. Notably, all outcomes had very low case‒control effect sizes, which might due to the employed group-level analysis methods, in which high heterogeneity would conceal the potentially informative features. Additionally, those studies mainly focused on older children with ASD and age-related effects were rarely considered. Moreover, research on brain asymmetry in DD/ID is lacking. How individual brain asymmetry at the early life stage is linked to gene expression and clinical manifestations and whether the relevant multimodal cascade could differentiate ASD and DD/ID from each other have not yet been investigated.

To answer these questions, we analyzed a large structural MRI dataset comprising 1030 children under 8 years old, including 563 children diagnosed with ASD (472 with DD/ID and 212 without DD/ID), 36 children with DD/ID only, and 219 typically developing (TD) children. We first assessed the mean values and distributions of the atypical individual deviations in grey matter volume (GMV) asymmetry derived from the normative models for the ASD and DD/ID groups. Then, we employed canonical correlation analysis to explore the associations between asymmetry deviations and clinical and behavioural measures. Associations between atypical deviations in GMV asymmetry and transcriptional profiles were further explored with partial least squares (PLS) regression analysis and between-region similarity analysis. Gene annotations were interpreted through gene enrichment analysis (see the flowchart in Figure 1).

**Figure 1.**
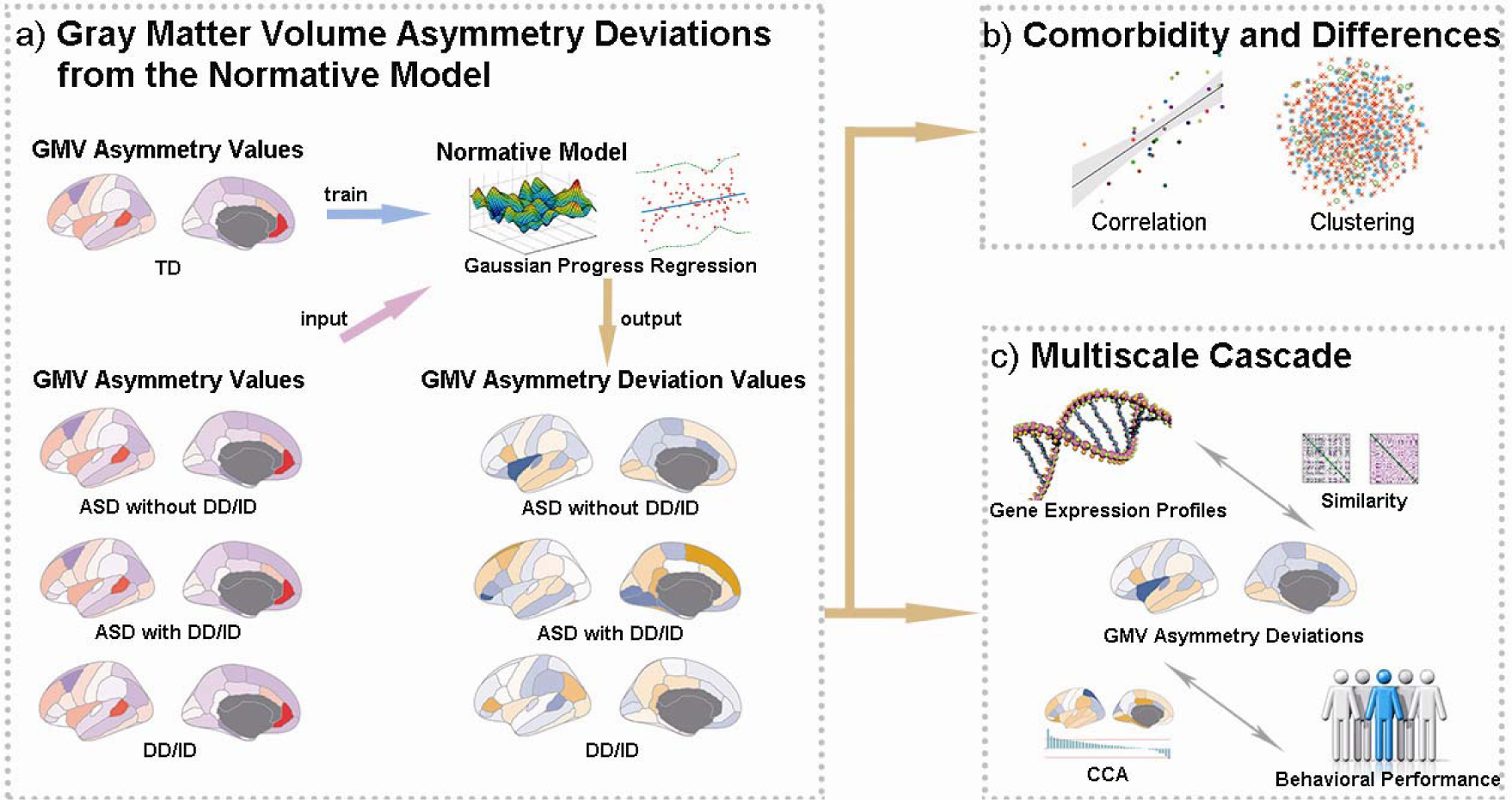
Overview: GMV asymmetry deviations in three atypical groups (ASD without DD, ASD with DD/ID, and DD/ID only) were extracted from a developmental normative model, and their associations with clinical symptoms and transcriptome profiles were calculated in a multimodal cascade involving a gene expression profile, grey matter volume (GMV) asymmetry deviations, and behavioural and cognitive measures. Abbreviations: ASD - autism spectrum disorder; DD/ID - developmental delay/intellectual disability; CCA - canonical correlation analysis.

## Results

### Sample characteristics

We included a large sample of 1030 young children from the Shanghai Autism Early Developmental (SAED) Cohort [40], including 563 children diagnosed with ASD with DD/ID (3.98 ± 1.22 years, range from 1.26 to 6.93 years, 472 males), 212 children diagnosed with ASD without DD/ID (3.24±1.15 years, range from 1.13 to 6.95 years, 184 males), 36 children with DD/ID only (4.42±1.4 years, range from 1.26 to 6.8 years, 25 males) and 219 age-matched typically developing children (4.42±1.62 years, range from 1.17 to 7 years, 107 males) as shown in Supplementary Table 1. The clinical measurements, including scores of Autism Diagnostic Observation Schedule (ADOS), Childhood Autism Rating Scale (CARS), Autism Behavior Checklist (ABC), the Infant-Junior High School Life Ability Scale (SM), and Social Responsiveness Scale (SRS), and cognitive performance, scores of Gesell Developmental Schedules (GDS) or IQ measured by Wechsler Preschool and Primary Scale of Intelligence (WPPSI) and Wechsler Intelligence Scale for Children (WIS-R), were statistically compared across the 3 diagnosis-related groups and the TD group. Specifically, we found the following: 1) the GDS or IQ scores of all 3 diagnosis groups are significantly lower than those of the TD group; 2) the GDS or IQ scores of ASD with DD/ID and DD/ID children are also significantly lower than those of ASD without DD/ID children; and 3) the two ASD groups had more severe social deficits than the TD and DD/ID groups (Bonferroni corrected, *P*s < 1×10^-8^). More details can be found in the Methods section and Supplementary Table 2.

### Atypical GMV asymmetry deviation patterns

We calculated the structural asymmetry metric employing the Desikan–Killiany atlas [41] for all subjects. The group averaged GMV asymmetry maps for the TD and 3 atypical groups are shown in Figure S1. Consistent with previous findings, similar structural lateralization patterns were found across 3 diagnosis-related groups and TD children. The superior temporal sulcus, rostral anterior cingulate and insula showed leftwards laterality. The caudal middle frontal cortex, parahippocampal cortex, prefrontal cortex and temporal pole showed rightwards laterality.

To determine the individualized GMV asymmetry deviation value of each region for each subject, we calculated sex-specific normative age models with the Gaussian process regression (GPR) method. Specifically, we trained the GPR models of GMV asymmetry for each region of interest (ROI) based on 247 TD children with age and sex as covariates. Then, we put the GMV asymmetry and covariate variables (age and sex) of children in ASD with DD/ID, ASD without DD/ID and DD/ID only groups into the generated models and obtained individual deviation values from the distributions from TD children. For each atypical group, regions with significant abnormalities in deviation values resulting from the one-sample t test are shown in Figure 2a (FDR corrected *p* < 0.05, Supplementary Table 3). Specifically, for ASD children without DD/ID, the inferior parietal cortex and precentral cortex showed significant negative deviations, indicating more rightwards asymmetry compared with that of TD children, while the rostral middle frontal cortex showed significant positive deviations, indicating more leftwards asymmetry (FDR corrected *p* < 0.05). Significant negative deviations in the inferior parietal cortex and precentral cortex were also observed in ASD children with DD/ID. Moreover, ASD children with DD/ID exhibited six additional ROIs showing significant atypical asymmetry, including the fusiform and entorhinal, which had more rightwards asymmetry, and the isthmus cingulate, the bank of superior temporal sulcus, the paracentral gyrus, and the rostral anterior cingulate cortex, which had more leftwards asymmetry. Notably, no significant deviations were found in any ROIs in children with DD/ID.

**Figure 2.**
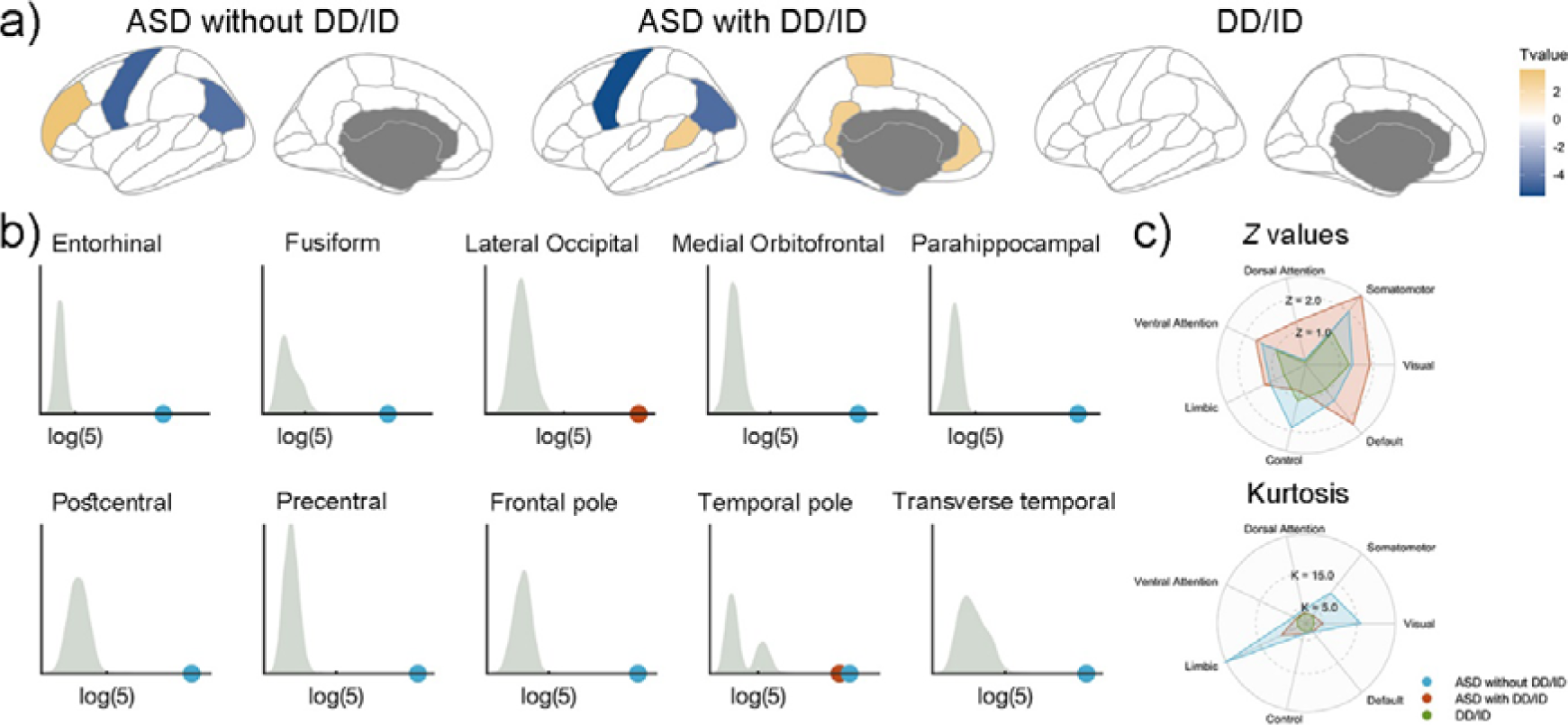
The GMV asymmetry deviations in the three diagnosis groups. **a).** The *brain regions with* significantly altered GMV asymmetry deviations characterized by one-sample *t* test in the three diagnosis groups. **b).** The brain regions with abnormal GMV asymmetry deviation distribution in ASD patients with and without DD/ID. The grey areas of the histograms show the kurtosis distribution of GMV deviations in TD children. The red and blue dots show abnormal kurtosis in ASD children with DD/ID and ASD children without DD/ID, respectively. **c).** The *average Z values* and kurtosis in Yeo’s 7 networks. The *t* value in each brain region was transformed into a Z value and used to calculate the average Z value for each network. Red indicates ASD children with DD/ID, blue indicates ASD children without DD/ID, and green corresponds to children with DD/ID only. **Abbreviations: ASD - autism spectrum disorder; DD/ID - developmental delay/intellectual disability.**

In addition to testing mean values, the kurtosis of the regional deviation values across subjects within each diagnosis group was calculated to explore whether the altered distribution differed from a normal distribution. Given that the kurtosis of the normal distribution is 3, a value above the threshold of 5 indicated an abnormality threshold in the current study. As shown in Figure 2b, the ASD without DD/ID group exhibited up to 11 regions showing extreme kurtosis (entorhinal, kurtosis = 64.391; fusiform, kurtosis = 31.153; medial orbitofrontal, kurtosis = 21.533; parahippocampal cortex, kurtosis = 50.502; postcentral cortex, kurtosis = 18.795; precentral cortex, kurtosis = 16.193; frontal pole, kurtosis = 23.538; temporal pole, kurtosis = 41.451; transverse temporal, kurtosis =16.145), indicating that the distribution of GMV asymmetry deviations was widely altered with high individual variations across subjects within the group. For ASD children with DD/ID, the lateral occipital (kurtosis = 12.203) and temporal pole (kurtosis = 32.759) showed a changed distribution of GMV asymmetry deviations. Again, no regions in the DD/ID group showed altered kurtosis values greater than the threshold (kurtosis < 5).

To characterize the extent and distribution of structural asymmetry deviations at the systems level, the mean Z values (normalized t values) and kurtosis values in Yeo’s 7 network [42] were calculated, as shown in Figure 2c. Similar ASD common atypical network deviation patterns were found in the default mode network, ventral attention network, somatomotor network and visual network. Atypical distribution patterns were also observed in ASD children with and without DD/ID with extreme heterogeneity in limbic and visual networks.

To explore the between-group differences, we conducted the two-sample *t*-test between every pair of diagnosis groups (ASD with DD/ID vs. ASD without DD/ID, ASD with DD/ID vs. DD/ID, and ASD without DD/ID vs. DD/ID) and found no significant between-group differences for any region after FDR correction (*p* < 0.05). Additionally, we found significant spatial correlations of GMV asymmetry deviations between the ASD with DD/ID vs. ASD without DD/ID groups (r = 0.557, permuted *p* = 0.0007), and between the ASD with DD/ID vs. DD/ID groups (*r* = 0.741, permuted *p* < 0.0001), as shown in Figure 3a.

**Figure 3.**
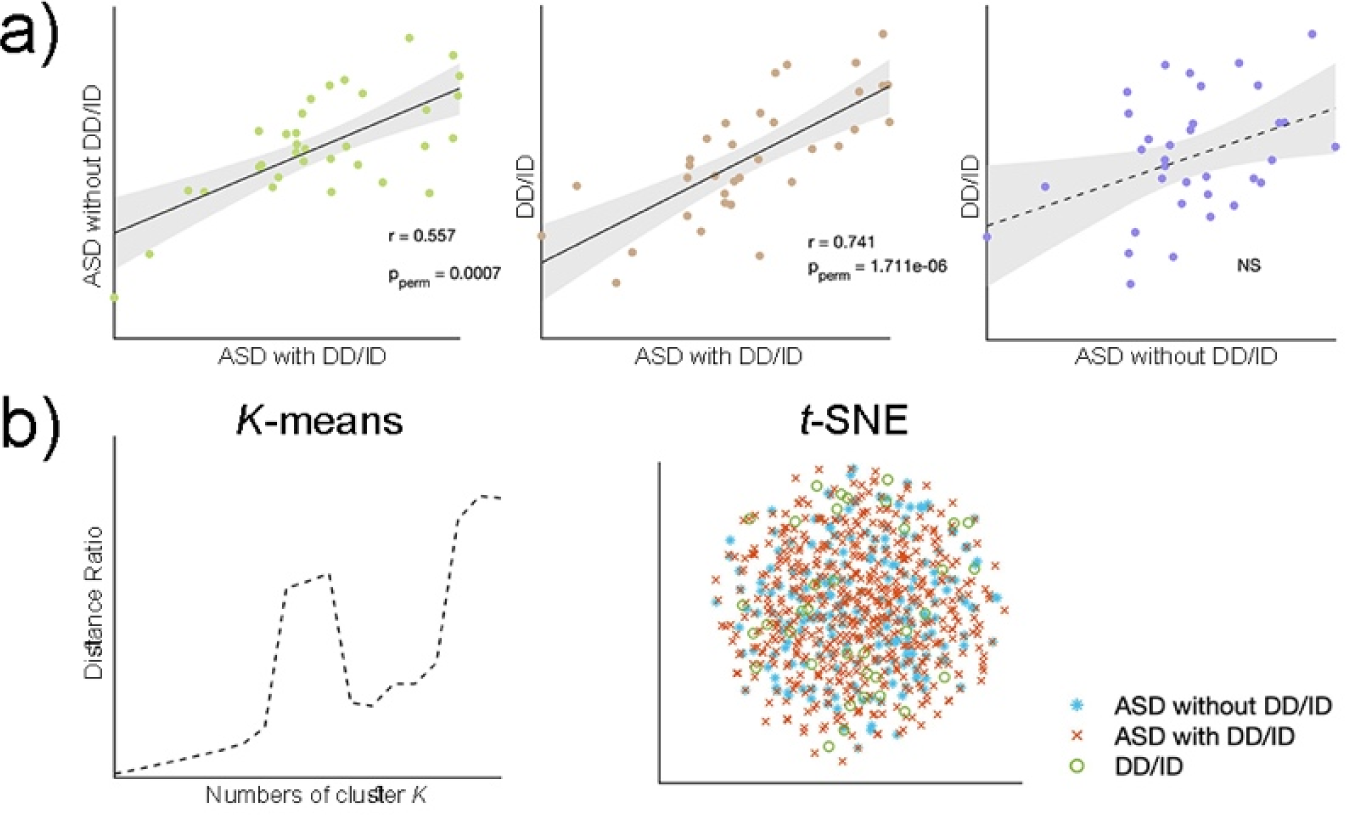
Comorbidities between diagnosis groups. **a).** Spearman correlations of whole-brain GMV asymmetry deviations between any two diagnosis groups. Significantly similar GMV asymmetry patterns were found in ASD with DD/ID v.s. ASD without DD/ID groups and ASD with DD/ID vs. DD/ID groups. **b).** The results of neither k-means clustering nor t-SNE can distinguish the 3 diagnosis groups from each other. **Abbreviations: ASD - autism spectrum disorder; DD/ID - developmental delay/intellectual disability.**

Through unsupervised clustering algorithms, we also found that these diagnosis groups cannot be separated. Through the *k*-means method, we calculated the distance ratio, as summed distances between-clusters divided by the summed distances within clusters, with the number of clusters from 2 to 20 and found no peak point as shown in Figure 3.b, indicating children with different diagnosis cannot distinguished by multivariate GMV asymmetry deviations. The spatial visualization by individual two-dimensional vectors reduced from 34-dimensional vectors via *t*-SNE also confirmed that children in the groups ASD with DD/ID, ASD without DD/ID and DD cannot be differentiated from each other.

### Associations between atypical asymmetry deviations and clinical symptoms

To understand the multimodal cascade in both the ASD and DD/ID populations, we conducted a canonical correlation analysis to explore the associations between brain structural asymmetry deviations and behavioural scores for ASD children without DD/ID, and ASD children with DD/ID respectively. For ASD children without DD/ID, GMV asymmetry deviations showed significant multivariate associations with clinical scores in one mode (*r =* 0.977, *p*_perm_ = 0.033). Six ROIs (paracentral, loading = 0.361; parahippocampal, loading = 0.295; inferior frontal gyrus pars opercularis, loading = 0.269; entorhinal, loading = 0.261; frontal pole, loading = 0.211; precentral, loading = 0.201) with positive loadings and four ROIs (caudal anterior cingulate, loading = −0.376; inferior temporal gyrus, loading = −0.293; rostral middle frontal gyrus, loading = −0.255; cuneus, loading = −0.201) with negative loadings contributed to the identified canonical mode (threshold of loading = ±0.2). Visual responsiveness (loading = 0.444), ADOS_2 (loading = 0.278), the sum of ADOS_1 and ADOS_2 (loading = 0.247), verbal communication (loading = 0.227), interaction (loading = 0.214), and independent living (loading = 0.212) scores showed positive contributions while intellectual consistency showed negative contributions (loading = −0.229) to the mode, as shown in Figure 4. Here, positive canonical loading indicates increasing neuropsychological scores with leftwards GMV asymmetry deviations.

**Figure 4.**
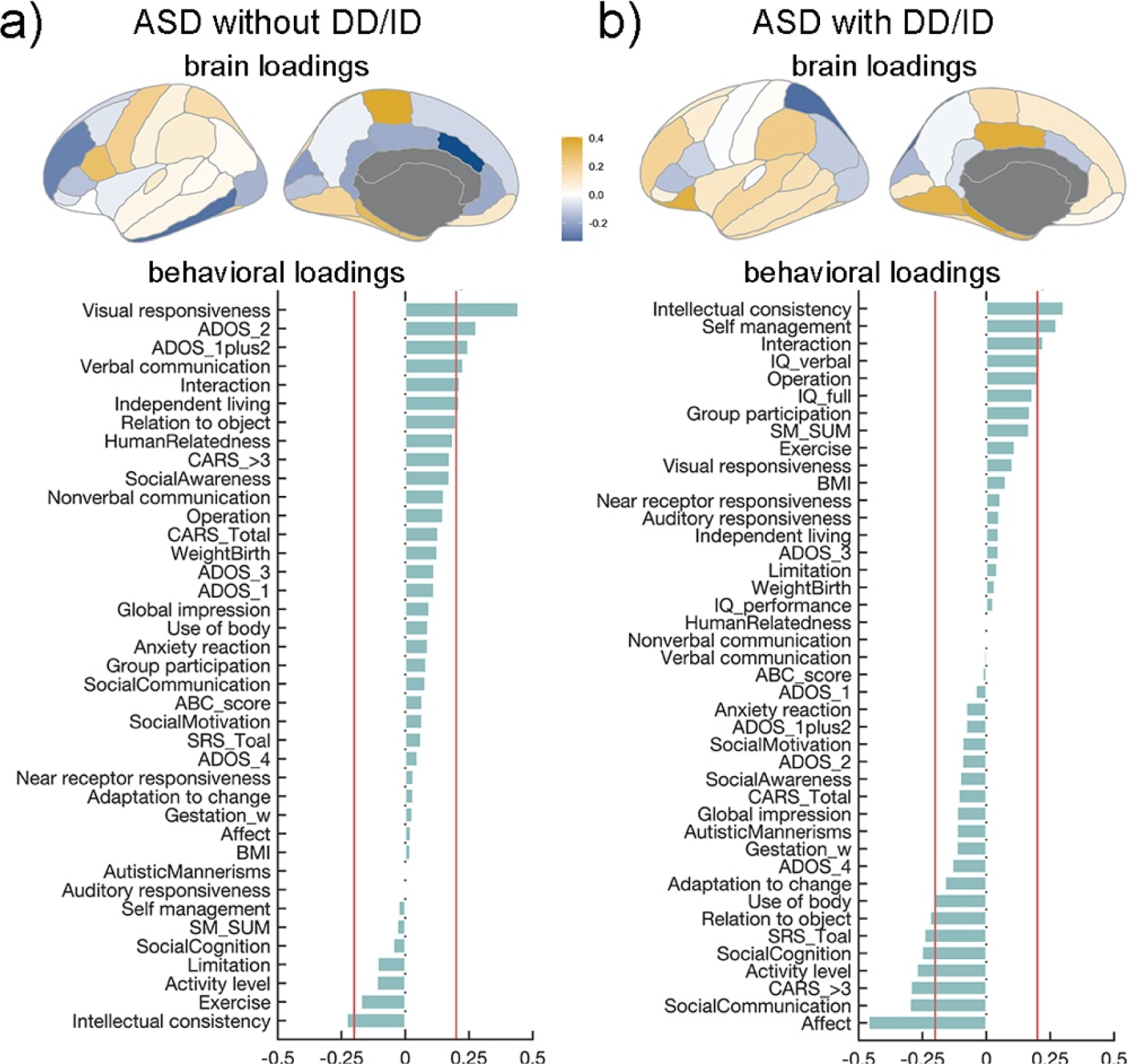
The identified canonical modes linking GMV asymmetry deviations and behavioural performance in ASD children without DD/ID and ASD children with DD/ID. **a).** The loadings of brain regions and ASD symptoms contributing to the canonical mode in ASD children without DD/ID. **b).** The loadings of brain regions and ASD symptoms plus IQ contributing to the canonical mode in ASD children with DD/ID.

For ASD children with DD/ID, GMV asymmetry deviations were identified in one significant canonical mode (*r =* 0.988, *p*_perm_ = 0.003) linking clinical scores and IQ scores. Eight ROIs (parahippocampal, loading = 0.405; posterior cingulate, loading = 0.369; lateral orbitofrontal, loading = 0.362; lingual, loading = 0.337; entorhinal, loading = 0.328; temporal pole, loading = 0.275; supramarginal, loading = 0.228; frontal pole, loading = 0.2) positively contributed to the identified mode. The superior parietal cortex was found to negatively contribute to the mode (loading = −0.325). In the linked clinical and IQ performance, intellectual consistency (loading = 0.303), self-management (loading = 0.247), interaction (loading = 0.222), verbal IQ (loading = 0.209) and Operation (loading = 0.2) showed positive contributions, while affect (loading = −0.46), social communication (loading = −0.299), CARS (loading = −0.295), activity level (loading = −0.272), social cognition (loading = −0.252), SRS (loading = −0.243), relation to object (loading = −0.22) and the use of body (loading = −0.2) showed negative contributions.

The canonical correlation analysis for ASD children with and without DD/ID did not yield any significant modes.

### Association between atypical asymmetry deviations and gene expression profiles

To quantify the multivariate association pattern between genes and the brain, we conducted the partial least squares regression and identified the regression components between gene expression profiles and GMV asymmetry deviations for each group. For the ASD with DD/ID group, regression component one explained 20.6% of the response variables (*r* = 0.454, *p* = 0.007), regression component two explained 14.7% of the response variables (*r* = 0.384, *p* = 0.025) and regression component three explained 16% of the response variables (*r* = 0.4, *p* = 0.019). For ASD without DD/ID group, regression component one did not significantly explain the response variables (*r* = 0.335, *p* = 0.053), regression component two explained 26.4% of the response variables (*r* = 0.39, *p* = 0.023), and regression component three explained 42.6% of the response variables (*r* = 0.402, *p* = 0.018). However, none of these correlations survived the spatial permutation tests (N = 10000, all *p*_*perm*^*s*^_ >0.5). Given that genes served as source factors, these results suggested that the association pattern between the gene expression profile and GMV asymmetry was not linear.

Next, inter-regional similarity analysis was conducted to detect significant associations between gene expression profiles and GMV asymmetry deviations for each diagnosis group, as shown in Figure 5a (for ASD children with DD/ID, r = 0.16, *p*_*perm*_ = 0.0001; for ASD children without DD/ID, r = 0.216, *p*_*perm*_ = 0; for DD/ID children, r = 0.241, *p*_*perm*_ = 0). The GCI values of autism related genes and intellectual disability related genes according to the In Situ Hybridization gene lists were extracted. A significant negative correlation was found between the GCI values of intellectual disability related genes between the DD/ID and ASD without DD/ID groups (r = −0.638, p < 0.001), as shown in Figure 5b.

**Figure 5.**
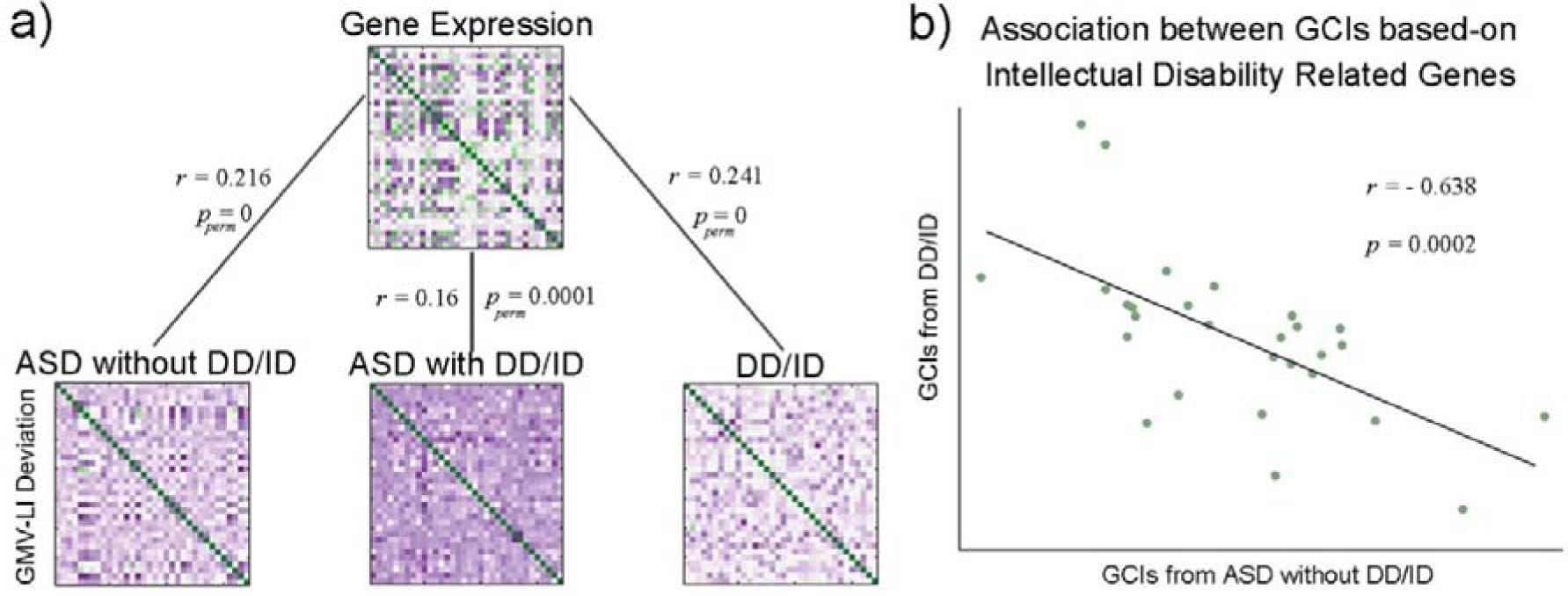
Associations characterized by interregional similarities between gene expression profiles and GMV asymmetry deviations of diagnosis groups. **a).** Significant interregional similarities were found between transcriptional profiles and GMV asymmetry in each group. **b).** The GCI values of intellectual disability-related genes were significantly negatively correlated between children with DD/ID only and ASD children without DD/ID. **Abbreviations: ASD - autism spectrum disorder; DD/ID - developmental delay/intellectual disability; GCI – gene contribution index.**

The shared and specific genes and their terms from the 3 diagnosis groups are shown in a circle network in Figure 6.a. For multigene list meta-analysis, the functional terms or pathways provide more information as shown in Figure 6b. We found four Gene Ontology (GO) terms of GMV asymmetry-related genes (‘regulation of transmembrane transport’, ‘neuron projection development’, ‘axon’, ‘supramolecular fiber organization’), that were consistent across ASD with DD/ID, ASD without DD/ID and DD/ID groups with FDR corrected *p* < 0.05. The GO term clusters specifically identified in ASD children with and without DD/ID were ‘postsynapse’ and ‘brain development’. The GO terms ‘cytoplasmic translation’ and ‘polymeric cytoskeletal fiber’ were clustered together in the ASD with DD/ID and DD/ID groups. Furthermore, ASD without DD/ID group exhibited the most of specific Go terms, including ‘nuclear inner membrane’, ‘synapse organization’, ‘regulation of synapse pruning’, ‘response to salt’, ‘cytoplasmic side of membrane’, ‘kinase binding’ and ‘perinuclear region of cytoplasm’, that were enriched for expression of specific genes associated with GMV asymmetry. GO terms of ASD with DD/ID specific genes were ‘regulation of neurotransmitter levels’, while that of DD/ID were ‘cell adhesion molecule binding’, ‘cell projection assembly’, and ‘regionalization’. To illustrate these findings, Figure 6c shows the proportion of each diagnosis group relating to the enriched GO term clusters, and Figure 6d shows the GO terms colored by cluster.

**Figure 6.**
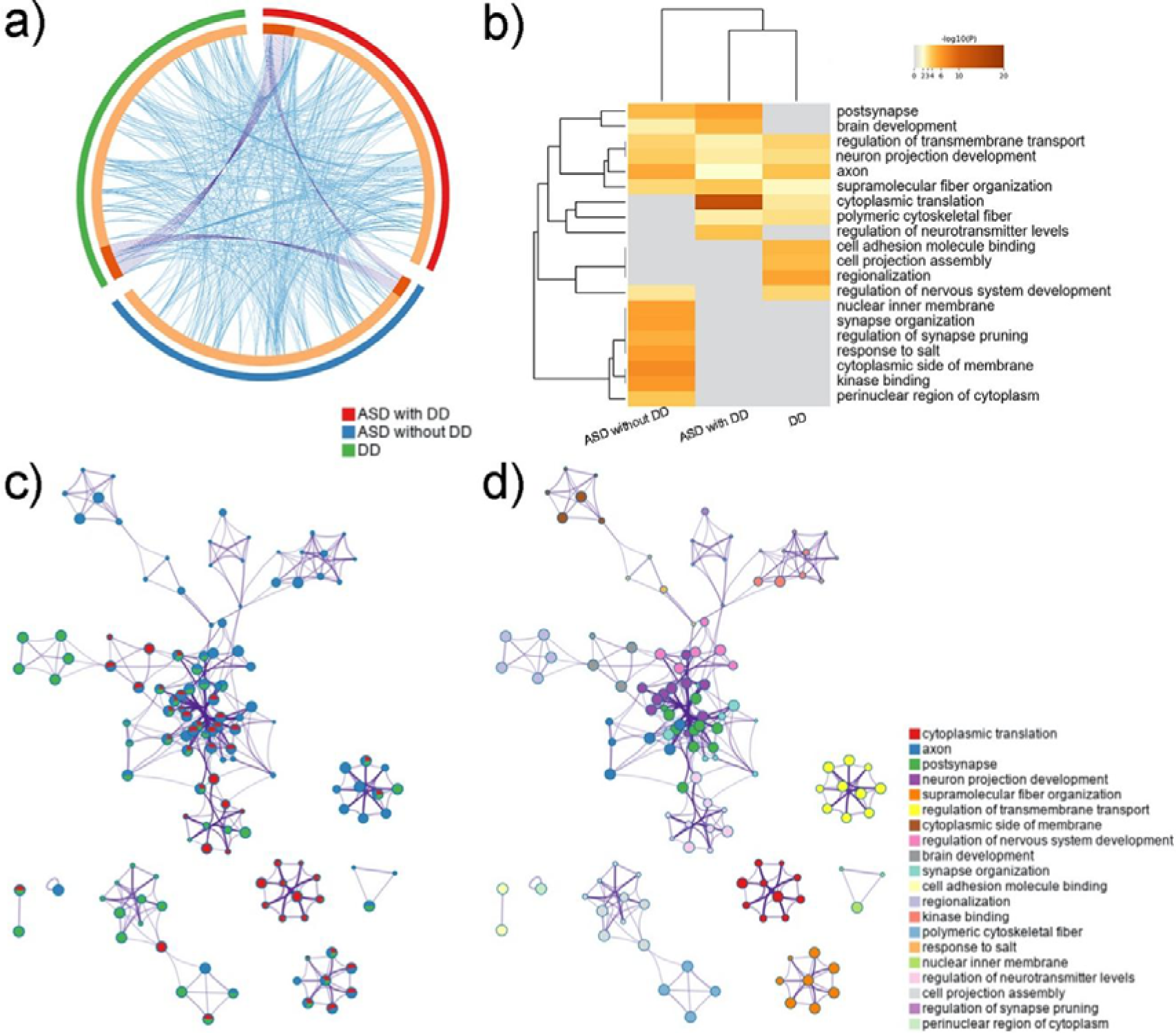
Functional enrichment of GMV asymmetry-related multiple gene lists from ASD children with DD, ASD children without DD and children with DD only. **a).** The shared and specific genes and their GO terms among diagnosis groups. Orange represents genes in each diagnosis group. Purple curves indicate identical genes between the two groups. Blue links show the same terms in which genes from the two groups were enriched. **b).** The top 20 clusters for enriched GO terms from all diagnosis groups, corrected by FDR *p* < 0.05. **c).** The GO term network characterized by pie charts. One circle represents an enriched term. The proportional coloured pie shows the percentage of genes from one diagnosis group in all groups; red indicates ASD children with DD/ID, blue represents ASD children without DD/ID and green indicates children with DD/ID only. **d).** The GO term network characterized by clusters. The same as c), but the colour of the circles indicates clusters. **Abbreviations**: **ASD - autism spectrum disorder; DD/ID - developmental delay/intellectual disability; GO – Gene Ontology.**

## Discussion

Using a sex-specific developmental normative model and GMV asymmetry as an intermediate variable, this study explored the neuropathological mechanisms underlying the comorbidities and differentiations between ASD and DD/ID in young children. We observed the ASD-specific GMV rightwards laterality in the inferior parietal cortex and precentral cortex, as well as abnormal heterogeneity in the temporal pole. Specifically, ASD children with DD/ID exhibited more regional abnormalities; ASD children without DD/ID showed higher within-group variability, while children with DD/ID showed no significant abnormalities. The GMV laterality of ASD children without DD/ID was associated with ASD symptoms, whereas that of ASD children with DD/ID was associated with both ASD symptoms and verbal IQ. Last, the GMV laterality of ASD children without DD/ID, ASD children with DD/ID, and children with DD/ID was associated with shared and unique gene expression profiles. However, those associations with intellectual genes in the ASD without DD/ID and DD/ID groups showed opposite effects.

Employed normative models, we detected significant regional abnormalities in GMV asymmetry in the inferior parietal cortex and precentral gyrus with more rightwards laterality and in the temporal pole with increased heterogeneity across subjects for ASD children both with and without DD/ID. The cognitive functions supported by these brain regions were core deficits in ASD populations, with the inferior parietal cortex involved in visuospatial processing [43, 44], the precentral cortex supporting sensorimotor functions, and the temporal pole involved in individual differences in sensory processing [45, 46], semantic representations [47], and social and emotional cognition [48]. This provides evidence that the current study captures the specific alterations of brain structure laterality in young children with ASD. In contrast to previous studies reporting inconsistent or non-regional specificity of GMV asymmetry in ASD patients [39, 49, 50], the current study calculated GMV asymmetry based on deviations from a normative model, which has shown potential in evaluating individual performance and understanding disease heterogeneity [51–54]. Additionally, the focus on young children in the current study minimizes the effects of developmental factors and their complex interactions with disease factors, suggesting that alterations in brain structure are only correlated with ASD. Notably, no significant deviations in GMV asymmetry were found in children with DD/ID. One hypothesis is that DD/ID involves impairments in multiple cognitive processes, including both the left and right brain systems, with little overall asymmetry [54].

The GMV asymmetry deviations of ASD children with DD/ID showed high overall similarities between ASD children without DD/ID and children with DD/ID only, whereas no significant correlation was found between the latter two groups. These results collectively indicated that the shared brain structural abnormalities between ASD children with DD/ID and without DD/ID are different from those between ASD children with DD/ID and children with DD/ID only. A reasonable hypothesis is that the former may point to the structural variants related to ASD, while the latter suggests the brain morphological abnormality linked to DD/ID. Subsequent studies could further explore this hypothesis by comparing other brain measurements with a normative model. In line with the previous work of Aglinskas et al., [55] showing non-subtype with brain structural features from ASD sources, this study found further evidence of whole-brain structural asymmetry similarity between the ASD with DD/ID and ASD without DD/ID groups, as well as between the ASD with DD/ID and DD/ID groups by unsupervised clustering. Echoing the subtle variants revealed by the normative model and overall similarity by correlation analysis, they collectively indicated that the brain GMV asymmetry deviations in ASD without DD/ID, ASD with DD/ID, and DD/ID only groups are gradually differentiated in a continuum, with ASD with DD/ID located in the middle.

The canonical correlation analysis identified a multivariate correlation mode between brain GMV asymmetry deviation and autistic symptoms in the ASD without DD/ID group, whereas those in the ASD with DD/ID group were associated with both ASD symptoms and verbal IQ. With more abnormal asymmetry in the parahippocampal, entorhinal, and frontal pole, involved in memory, exploiting and exploration [48], both ASD children with and without DD/ID showed more serious communication and social impairments. Congruent with this, similar abnormal GMV asymmetry of the above brain regions and their links to social communication problems have been reported in previous studies [56–59]. Of note, canonical correlation analysis failed to identify brain GMV asymmetry-clinical phenotype associations in all ASD children. This may have been due to the high intragroup heterogeneity combining brain GMV asymmetry abnormalities from common ASD effects and DD/ID effects, which yielded a complex relationship between brain structural changes and clinical symptoms and cannot be identified by linear strategies in canonical correlation analysis. Future studies could further explore this by investigating the heterogeneity of ASD, or using nonlinear algorithms to establish the links between brain measurements and clinical phenotypes.

The shared and specific genes and their enriched terms were identified for the 3 diagnosis-related groups while ASD without DD/ID group exhibits the most ones. GMV asymmetry related genes were enriched in the GO terms ‘postsynapse’ and ‘brain development’ in ASD children with and without DD/ID. This suggested that the above gene functional abnormalities may be microscale characteristics of ASD; an extensive number of studies have supported this finding [60, 61]. Our results also showed that the GO terms ‘cytoplasmic translation’ and ‘polymeric cytoskeletal fibre’ were associated with ASD with DD/ID and DD/ID alone. Given the overall similarity of whole-brain GMV asymmetry between ASD children with DD and those with DD only, we suggest that cellular dysfunctions affect brain morphology and further drive complex intellectual delay phenotypes.

Here, we address two unique strengths in this study One is the unique ASD data on young children ranging in age from 1-7 years from the SAED cohort, which has a large sample. The other is the combination of a normative model and multimodal cascade. The former allows the extraction of individual abnormalities, and the latter enables a comprehensive perspective. The study also had limitations. First, the covariates of the normative model extracting GMV asymmetry deviations were age and sex, but environmental factors of family and community, such as socioeconomic status, may also be related to brain morphology development [62, 63]. These potential factors could be included in future studies. Second, limited by the availability of gene expression data, we linked GMV asymmetry deviations in 1- to 7-year-old children to transcriptional profiles from healthy adults, which may have affected the results. With the gradual establishment and improvement of ASD datasets, follow-up analysis is expected to investigate the brain morphology-transcriptome association within the same ASD participants. Finally, functional brain variants play an important role in the multimodal cascade. How functional brain metrics may mediate the relation between the structural brain and the behavioural phenotype is worthy of further exploration. The hierarchical associations of gene expression-brain morphology-brain functions-clinical phenotypes may elucidate the deeper pathological mechanism of ASD heterogeneity and DD.

In conclusion, our study demonstrated the transcriptome-brain structure-behaviour cascade, as well as their comorbidities and differences, in children with ASD and DD/ID under 8 years of age from the SAED cohort. The brain GVM asymmetry abnormalities in ASD children without DD/ID, ASD children with DD/ID, and children with DD/ID only were similar overall, but there were subtle differentiation patterns that are linked to diagnosis-specific clinical phenotypes as well as transcriptome profiles. Of note, ASD-specific GMV showed rightwards laterality in the inferior parietal cortex and precentral cortex, as well as abnormal heterogeneity in the temporal pole. Moreover, GMV laterality, as a central feature of autism, is linked to autistic traits and cognitive performances, with marked deviations from controls in terms of development. Future work may explore the potential applications of brain asymmetry for ASD subtyping and clinical screening.

## Methods

### Participants

Participants were recruited in the neurodevelopmental project - Shanghai Autism Early Development Cohort[40]. All children were recruited from those who had visited the Department of Developmental Behaviour and Children Health Care, Xinhua Hospital, Shanghai Jiao Tong University, due to concerns of linguistic and/or social deficits and were assessed for early signs of autism. This study employed structural MRI data from 1030 young children. After imaging quality control (see below for details), the data of 1030 young children were employed for analysis, including 563 children diagnosed with ASD and DD/ID (3.98±1.22 years, range from 1.26 to 6.93 years, 472 males), 212 children diagnosed with ASD without DD/ID (3.24±1.15 years, range from 1.13 to 6.95 years, 184 males), 36 children with DD/ID only (4.42±1.4 years, range from 1.26 to 6.8 years, 25 males) and 219 age-matched TD children (4.42±1.62 years, range from 1.17 to 7 years, 107 males). The clinical diagnoses of ASD were made by pediatricians according to the *Diagnostic and Statistical Mental Disorders, Fifth Edition* (DSM-5) and ADOS or ADOS-toddler. To determine whether a child has DD/ID, his or her general intellectual ability was assessed by the GDS for those younger than 4 years, WPPSI for those aged 4–6 years and the Wechsler Intelligence Scale for Children (WIS-R) aged 6–7 years, with the GDS cut-off score < 75 or a full IQ < 70. The study was approved by the Ethics Committee of Xinhua Hospital affiliated with the Shanghai Jiao Tong University School of Medicine (XHEC-C-2019-076) and registered with ClinicalTrials.gov (NCT04358744).

All patients completed ASD-related neurocognitive assessments, including the ADOS, CARS, ABC, and SRS. The total scores and subitem scores were used for subsequent analysis.

### MRI data acquisition, quality control and processing

Participants received MRI scans on one of two MRI scanners: a Siemens Verio or a Philips Ingenia 3.0T MRI scanner. Similar T1-weighted axial protocols were employed for structural MRI data collection. For the Siemens Verio scanner, the following parameters were used: repetition time (TR) = 2300 ms, echo time (TE) = 2.28 ms, flip angle = 8°, FOV = 192 mm * 192 mm; voxel size = 1*1*1 mm^3^; slice thickness = 1 mm, and slice number = 170. For the Philip Ingenia scanner, the following parameters were used: TR = 7.9 ms, TE = 3.5 ms, flip angle = 8°, FOV = 250 mm * 199 mm; voxel size = 1*1*1 mm^3^; slice thickness = 1 mm, and slice number = 170. To ensure collection, uncooperative children were sedated with chloral hydrate orally or by enema. Highly cooperative children were scanned in a resting state. The whole scanning procedure was conducted under the supervision of medical staff.

Raw structural images were independently manually inspected by two staff members (Dr. Y Dai and M.D. Y Liu) to exclude those with obvious motion artefacts. After quality control, the preprocessing steps included skull stripping, and realignment using CAT12 and then reregistered and segmented based on the Chinese pediatric atlases (CHN-PD) [64]. Then, we conducted the second round of quality control, comprising algorithms and a manual inspection of the segmentation results. Finally, smoothing using SPM12 with an 8 mm isotropic Gaussian kernel was conducted.

After preprocessing, we calculated the structural asymmetry metric employing the Desikan–Killiany atlas [41]. First, we coregistered the atlas to the CHN-PD and parcellated the preprocessed structural images of all subjects based on it. Then, the GMV for each brain region was calculated as the sum of voxelwise intensities. Finally, the laterality index in each region for every subject was calculated as 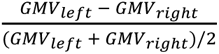, which is a commonly used formula for values with a range from −2 to 2. The ComBat analysis based on an empirical Bayes approach was utilized to harmonize the site effects [65–67].

### Statistical and normative modelling analysis

To generate the individualized abnormal asymmetry profiles of each region, we calculated the sex-specific normative age models using the Predictive Clinical Neuroscience toolkit [68] with the Gaussian process regression (GPR) method, which can provide point estimates as well as coherent measures of predictive confidence. Specifically, GPR models of GMV asymmetry for each ROI were trained based on 247 TDs with age and sex as covariates. Fivefold cross-validation was conducted to avoid overfitting and obtain optimal parameters. In this way, we generated developmental normative GMV asymmetry distributions in TD individuals. Then, we put the individual GMV asymmetry metric and covariate variables (age and sex) of children in the ASD with DD/ID, ASD without DD/ID and DD/ID only groups into the generated models and obtained the individual Z scores, which indicated the statistical estimate of the degree of deviation from TD distributions for each ROI of each subject.

To depict the group-specific profiles, we calculated the extents and distributions of asymmetry deviations for each atypical group. First, a one-sample *t* test was performed on deviation values for each group with a false discovery rate correction (FDR *q* < 0.05). Notably, a positive *t* value indicated that the corresponding ROI was more leftwards lateralized compared with that of TD children, while a negative *t* value indicated increasing rightwards asymmetry. In addition to testing the mean values, kurtosis of deviation values across subjects within each group for each region was calculated to explore whether the distribution alterations differed from a normal distribution. Considering that the kurtosis value of a standard normal distribution is 3, values above 5 were set as the threshold for abnormal criteria.

To investigate the categorical relationships among ASD and DD/ID populations, the similarities and differences in GMV asymmetry deviations among groups were then evaluated. Two-sample *t* tests were performed to capture between-group differences in mean values and kurtosis of deviation values of each region with FDR correction (FDR *q* < 0.05) for multiple comparisons. Between-group similarity was calculated through Spearman’s correlations across brain regions based on the group-averaged GMV asymmetry deviation values. To exclude the confounding effects of spatial autocorrelation [69, 70], spatial permutation analysis (n = 10,000 times) based on spherical rotations was conducted for each pair of diagnosis groups.

### Clustering

To further investigate the relationships between diagnosis groups, we conducted unsupervised clustering, *k*-means and *t*-distributed stochastic neighbour embedding (*t*-SNE) analyses using MATLAB internal functions. For *k*-means clustering, the summed between-group distances divided by the summed within-group distances were calculated as a function of cluster number *k*. The peak point of the generated ratio curve was selected as the optimal number of clusters, which indicated the highest homogeneity within groups with the lowest heterogeneity between groups. *t*-SNE is a nonlinear machine learning algorithm that converts high-dimensional data to a two-dimensional space for visualization. In this work, GMV asymmetry deviations in 34 regions for all subjects were imported into *t*-SNE to test whether subjects within the same diagnosis group were spatially gathered.

### Associations with clinical and neuropsychological measurements

To understand the multiscale cascade in ASD and DD/ID populations, we conducted a canonical correlation analysis to explore the associations between brain structural asymmetry deviations and behavioural scores for ASD children without DD/ID and ASD children with DD/ID separately and jointly. The clinical measurements included body mass index; gestational weight; weight at birth; ABC score; and the ADOS, CARS, SM, SRS total and sub-item scores. The neuropsychological measurements included the GDS scores or IQ scores. The clinical measurements and neuropsychological scores were put evaluated in canonical correlation analysis separately and jointly. Considering that age and sex were controlled as covariates in the normative model for GMV asymmetry deviation calculation, we did not include them here. The pairs of linear combinations of deviations in GMV asymmetry and clinical/neuropsychological measurements, known as canonical variables, were obtained. Then, we calculated the linear correlations of every original variable with the canonical variables, referred to as canonical loading, to determine their contributions. The statistical significance of the identified mode was tested through random permutation with 10,000 rows in the original matrix.

### Associations with gene expression profiles

#### Estimation of regional gene expression profiles

Here, we employed the genome expression data of six postmortem neuro-typical human brains from the Allen Human Brain Atlas (AHBA) dataset (http://human.brain-map.org) to explore the associations between deviations in GMV asymmetry and relevant gene expression profiles [71]. We only used the transcriptional data from the left hemisphere considering that the available right hemisphere data were from only two donors. Traditional pre-processing of AHBA data was conducted according to the procedures in Arnatkevic et al. [72]. Then, the regional gene expression matrix (34*10,027, expression levels of 10,027 genes across 34 brain regions) was obtained for further analysis.

#### Partial least squares regression

We conducted the partial least squares regression (PLS) to investigate the effects of transcriptome profiles on the group-specific deviation patterns in GMV asymmetry[73]. Specifically, for each group, the gene expression matrix was used as the predictor variables while the statistic GMV asymmetry deviation values (t values from the one-sample *t* test) were used as response variates. A spatial permutation (n = 10,000) based on spherical rotations [69, 70] was performed to test the significance of the covariance among the PLS components, i.e., the linear combinations of transcriptome levels and deviations in GMV asymmetry [74, 75]. Then, we used Pearson’s correlation to calculate the spatial similarity between the identified PLS components of gene expression and the GMV asymmetry deviation maps controlling the influences of spatial autocorrelation [69, 70]. Each gene’s contribution to the maximally covariant PLS components was represented as the corresponding PLS weights, whose variabilities and significances were estimated through bootstrapping (n = 10,000) [74, 76, 77]. In this way, we obtained a list of genes that would be altered corresponding to the deviation patterns in GMV asymmetry for each atypical group (p < 0.05, FDR correction, two-tailed).

#### Inter-regional similarity analysis

Inter-regional similarity analysis was also conducted to calculate to explore the relationship between genetic expression and GMV asymmetry patterns. Here, we utilized Pearson’s correlation to calculate the inter-regional similarity matrices (34 × 34) for gene expression profiles and deviations in GMV asymmetry separately. The lower triangle of the resultant matrices for each group was extracted as a vector. Then, the similarity between the vector of gene expression profiles and vector of deviations in GMV asymmetry was calculated with Spearman’s correlation analysis for each diagnosis group. Spatial permutation (n = 10,000) tests were conducted to exclude spatial autocorrelation effects.

To further quantify the contribution of each gene to the results of inter-regional similarity analysis, a leave-one-out procedure was conducted. Specifically, each gene was removed, and the inter-regional similarity analysis was constructed based on the remaining genes. The difference between the new Spearman’s correlation coefficient and the original Spearman’s correlation coefficient was defined as the gene contribution index (GCI). To test the statistical significance of each GCI, bootstrapping (n = 10,000) with replacement of 10227 genes was performed to obtain the null distribution. The ratio characterized by the original GCI divided by the standard deviation of its null distribution was noted as the standard GCI score. Then, we ranked all genes by their standard GCI score and obtained an ordered, group-specific gene list.

#### Enrichment analysis

To explore the functional annotations of the gene lists for group-specific GMV asymmetry, we performed a meta-analysis for multiple gene lists based on the GO and the Kyoto Encyclopedia of Genes and Genomes (KEGG) on the Metascape website (https://metascape.org/gp.inedx.html#/main/step1). After extracting the top 1% of genes and the bottom 1% of genes from each ordered gene list, we uploaded the formed multiple gene list to the Metascape website, which has automated enrichment analysis tools with more than 40 independent knowledge bases. Then, the GO (biological processes, cellular components and molecular functions) terms and KEGG pathways enriched in each list were selected with FDR correction, *p* <0.05.

## Acknowledgements

This study was supported by grants from the National Natural Science Foundation of China (82125032, 81930095, 81901826 and 81761128035), the Science and Technology Commission of Shanghai Municipality (19410713500 and 2018SHZDZX01), the Shanghai Municipal Commission of Health and Family Planning (GWV-10.1-XK07, 2020CXJQ01, 2018YJRC03), the Shanghai Clinical Key Subject Construction Project (shslczdzk02902), the Shanghai Municipal Science and Technology Major Project [No.2018SHZDZX01], Innovative research team of high-level local universities in Shanghai (SHSMU-ZDCX20211100), the Guangdong Key Project (2018B030335001), the National Nature Science Foundation of China (82204048), the National Natural Science Foundation of China (82001771) the Shanghai Municipal Commission of Health and Family Planning (20214Y0125).

## Author contributions

M.C. and F.L. designed the study. Y.L., Y.D., L.D., Z.C., Y.Z., M.T., L.Z., T.R. contributed to the acquisition of research data. S.G., and M.C. conducted the data analysis. F.L., M.C., and S.G., provided the interpretation of results. S.G. and M.C. wrote the first draft of the manuscript. E.T.R., M.C., F.L. and J.F. revised the manuscript.

## Notes

### Competing Interest Statement

The authors have declared no competing interest.

